# Determination of the fixed oil quality of ripe *pistacia lentiscus* fruits and *Opuntia-ficus indica* seeds

**DOI:** 10.1101/2020.11.20.392084

**Authors:** Mohamed Kechidi, Mohamed Anis Chalal, Amel Bouzenad, Asma Gherib, Brahim Touahri, Mohamed Abou Mustapha, Mohamed Ourihene

**Affiliations:** Applied Biochemistry and Microbiology Laboratory, Department of Nature and Life Science, Benyoucef Benkhedda university (Algiers -1- university) Algiers, Algeria; Medical biology laboratory, Algerian pasteur institute annex El Hamma, Algiers, Algeria; REPI society (Agro-food and pharmaceutical industry) Beaulieu, Oued Smar, Algiers, Algeria (www.repi.dz); Center for scientific and technical research in physical and chemical analyses (CRAPC) Bou-Ismail, Tipaza, Algeria; Sabrinelle laboratory Bordj El Bahri, Algiers, Algeria

**Keywords:** Pistacia lentiscus, Opuntia-ficus indica, Lentisk oil, Prickly seeds oil, physicochemical parameters, fatty acids

## Abstract

*Pistacia lentiscus* and *Opuntia-ficus indica* are used in several fields, this study made it possible to highlight the determination of the oil quality from the fruits of *Pistacia lentiscus* and that of the seeds oil of *Opuntia-ficus indica*, and this, by determining its physicochemical parameters such as acid value, saponification and insaponification value, iodine index, peroxyd value, as well as refraction index, humidity and their biochemical compositions, in particular the fatty acids (by CPG) from the samples of oils collected from the region of Khemis Miliana (Ain defla) and extracted by a mechanical method. The results show a quantitative difference between the oily samples in percentage of fatty acids. The contents of Oleic Acid, Linoleic Acid and Palmitic Acid are highest in the case of lentisk oil and are respectively 58.35%, 19.65%, 19.63%. However, the content of Linoleic Acid, Oleic Acid, Palmitic Acid and Stearic Acid are highest in the case of prickly pear and are respectively 63.74%, 21.30%, 10.17%, 3.58%.

## 1. Introduction

*Pistacia lentiscus* (L) is a medicinal species. It is a shrub of the genus *Pistacia* belonging to the *Anacardiaceae* family **(Bozorgi et al., 2013)**, it appears as an evergreen tree or shrub with 2 to 3 meters high **(Alloune et al., 2012)**, grows throughout the Mediterranean basin in a subhumid, semi-arid and arid site of Europe, Africa and Asia, as far as the Canaries and Portugal **(Verdú and García-Fayos, 1998)** and it is commonly dispersed in Algeria over the entire littoral zone and along the tell and in forest areas **(More and White, 2005)**. This plant species is adapted to the stress resulting from the lack of water and is able to fight against erosion which is a key factor in the desertification of the ecosystem of semi-arid Mediterranean regions **(Dogan et al., 2003)**. *Pistacia lentiscus* oil is dark green in color, the *lentisk* fruit oil composition and its chemical characteristics were investigated. The studies show that Lentiscus oil consists mainly of unsaturated fatty acids (mono and polyunsaturated) and saturated fatty acids, accompanied by auxiliary lipid substances called minor constituents, such as tocopherols, phyto-sterols and phenolic compounds **(Dhifi et al., 2013).** In traditional medicine, the *Pistacia lentiscus* known for its analgesic, antibacterial, antifungal, antioxidant effect, It is used for the treatment of eczema and kidney stones and considered as an anticancer agent, in particular against tumors of the breast, liver, stomach, spleen, and uterus **(Assimopoulou and Papageorgiou, 2005).**

*Opuntia-ficus indica* known as prickly pear, a member of the *Cactaceae* family **(Mulas and Mulas, 2004).** It is a robust plant that can grow up to 5 meters in height with a thick, woody trunk **(Habibi, 2004).** Its geographical distribution is located mainly in: Mexico, Sicily, Chile, Brazil, Turkey, Korea, Argentina and North Africa **(Felker et al., 2005)**. In Algeria, prickly pear plantations are spread across the highlands **(Piedallu, 1935)**, It is planted for fruit consumption and also as an ornamental, for wind protection fencing, land reclamation and rehabilitation, and erosion control. *Opuntia-ficus indica* oil is light yellow to greenish in color. the differents studies show that the preakly pear seed oil is edible; it could be another nutritious and functional product of potential interest for the agro-industry, it is rich in unsaturated fatty acids **(Inglese, 2019)**. The *Opuntia-ficus indica* fruits is used as a natural anti-wrinkle agent and for the manufacture of anti-wrinkle skin creams **(Ennouri et al., 2005)** and it has been used for their hypoglycemic and hypolipidemic actions. Some authors have attributed these beneficial effects to high contents of fibers in these fruits **(Aires et al., 2004).** this oils are produced in Algeria, especially in the north of the country where the species abounds **(Leprieur, 1860)**.

The current study was undertaken to highlight the quality of *Pistacia lentiscus* and *Opuntia-ficus indica* fixed oils through their physicochemical characteristics and their compositions of fatty acids.

## 2. Material and methods

### 2.1. Material

#### Samples

*Pistacia lentiscus* and *Opuntia-ficus indica* ripe fruits were harvested from plants growing wild in in the region of Khemis Miliana wilaya of Ain Defla (North-ouest of Algeria) in december and august 2019 respectively

#### Fat Extraction

the extraction of fixed oil from the fruits of *Pistacia lentiscus* and the seeds of *Opuntia-ficus indica* was carried out by an industrial method known as: cold pressing extraction. In this process we used a REPI type screw press with a capacity of 50kg / h.

### 2.2 Methods

#### Determination of the physical properties of the oils

the refractive index of the two oils is determined by an ABBE refractometer, the water content and volatile matter is determined by an infrared dryer.

#### Determination of the chemical properties of the oils

the acid index and acidity, the saponification and unsaponification index, the peroxide index and the iodine index are determined respectively according to the standards protocol: ISO 660-1996, NA 276-1992, IUPAC 1987 2:401, ISO 3960-1977, ISO 3963-1989.

##### 2.2.3. Determination of the fatty acid profile of the two oils

###### Preparation of methyl fatty acid esters

Methyl esters were prepared according to the standard protocol AFNOR T60-233. we weighed 1g of oily sample and put it in a vial then added 10 ml of heptane then added 0.5 ml of the methanolic potassium solution we have already prepared before (11.2g of KOH in 100ml of methanol), we shake the solution for 20s and finally we collected the top layer containing the methyl esters.

###### GC-MS Analysis of Fatty Acids Methylic Esters (FAMS)

The chemical composition of vegetable oil was determined by a Gas phase chromatography such as Hewlett Packard Agilent 6890 plus coupled with Hewlett Packard Agilent 5973 mass spectrometry using HP-5MS capillary column with the following characteristics: length, 30 m; internal diameter, 0.25 mm; film thickness 0.25 μm. the carrier gas was the helium, at a flow through the column of 1 ml/min. Injection mode was SPLITLESS, the injector temperature was maintained at 250 C and the flame-ionisation detector (FID) at 230 C. The temperature of the oven has been programmed as follows: 2C/min from 60C to 280C / 8 min to 60C / 15 min to 280C. The injected volume of fixed oils is 0.1 μm. Each FAME present in the oil was identified by comparison of its retention time and mass spectrum with those of authentic compounds.

## 3. Results and discussion

Table 1 shows the determined amount of oils extracted from *Pistacia lentiscus fruits*, *Opuntia-ficus indica* seeds and their Physicochemical characteristics.

**Table 1:** the amount and the Physicochemical characteristics of *Pistacia lentiscus* and *Opuntia-ficus indica* oil

|  | *Pistacia lentiscus* | *Opuntia-ficus indica* |
| --- | --- | --- |
| Oil (%) | 20 | 3.25 |
| Refractive index | 1,463 | 1,468 |
| Moisture content (%) | 0.47 | 0.21 |
| Acid value (mg of KOH/g of oil) | 28,6 | 7,01 |
| Acidity (%) | 14,41 | 3,53 |
| Saponification value (mg of KOH/g of oil) | 269,28 | 222,3 |
| Unsaponification value (g/100g) | 1,6 | 1,8 |
| Peroxide value (meq O<sub>2</sub>/kg of oil) | 6,0 | 59,5 |
| Iodine value (g of I<sub>2</sub>/100 g of oil) | 44,6 | 45,8 |

The crude fat content of the *Pistacia lentiscus* fruit varied from 20%; The amount extracted from the *Opuntia ficus indica* was 3.25%.

**Table 2:** Fatty acid composition (percentage of the total fatty acids) of the *Pistacia lentiscus* and *Opuntia-ficus indica*.

|  | Oil of *Pistacia lentiscus* | Oil of *Opuntia-ficus indica* |
| --- | --- | --- |
| (C15:0) | - | 0,04 |
| (C16:0) | 19,63 | 10,17 |
| (C16:1) | 1,15 | 0,38 |
| (C18:0) | 1,00 | 3,58 |
| (C18:1) | 58,35 | 21,30 |
| (C18:2) | 19,65 | 63,74 |
| (C20:0) | 0,09 | 0,20 |
| (C20:1) | - | 0,27 |
| (C22:0) | 0,05 | 0,12 |
| (C22:1) | - | 0,11 |
| (C24:0) | 0,08 | 0,09 |
| SFA | 20,85 | 14,2 |
| UFA | 79,15 | 85,8 |
| U/S | 3,80 | 6,04 |

C number of carbon atoms in the fatty acid, (C15:0) Pentadecanoic acid, (C16:0) Palmitic acid, (C16:1) Palmitoleic acid, (C18:0) Stearic acid, (C18:1) Oleic acid, (C18:2) Linoleic acid, (C20:0) Arachidic acid, (C20:1) Gondoic acid, (C22:0) Behenic acid, (C22:1) Erucic acid, (C24:0) Lignoceric acid, SFA saturated fatty acids, UFA unsaturated fatty acids, U/S unsaturated/saturated fatty acids.

In the case of the ripe fruits of *Pistacia lentiscus*, our result is lower than that found by **(Djedaia, 2017)** about 64,5% and **(Trabelsi et al., 2012)** about 42,54%. However in the case of the seeds of *Opuntia-ficus indica*, our results is lower too than that found by **(Boukeloua, 2009)** about 10,45% and (**Salvo et al., 2002)** about 8-9%.

the yield of vegetable oil of the same or a different species may vary depending on several parameters namely the plant species, the harvest period and the extraction method.

The refraction index is a purity test **(Karleskind, 1992)**, it’s depends on the density, chemical composition of the oil and temperature. It increases with the *unsaturation and the presence of the fatty chains of secondary functions **(Boukeloua et al., 2012)**. The refractive index measured for the samples of our oils are from 1.463 for Pistacia lentiscus* and 1.468 for *Opuntia-ficus indica.* These values are similar to those reported by **(Karleskind, 1992)**, for olive, palm, and avocado oils, which are respectively (1,468-1,470) and (1,453-1,458) and (1,465-1,474), and in standards established by CODEX and FAO 2013. Which is from (1,4677-1,4705). This index allows us to classify our oils studied as non-siccative oils (1.467< RI > 1,472) **(Wolff, 1968)**.

*Pistacia lentiscus* oil contained 0,47% moisture and *Opuntia-ficus indica* oil contained 0,21%. Low water content is more than essential for oil stability, as water promotes alteration reactions **(Aïssi et al., 2009).**

The acidity is high in *Pistacia lentiscus* oil about 14.41% compared to the value cited by codex alimentarius (5%), while this index is low in the *Opuntia-ficus indica* oil about 3,53%. which indicates that Pistacia lentiscus oil contains a huge amount of free fatty acids. The high acidity values in the oils may be due to the poor preservation of the fruit before extraction and analysis and this may be explained by the hydrolysis of the triglycerides under the action of lipase contained in the fruits causing the release of free fatty acids or to Abnormalities during the biosynthesis process, microbial activities and environmental conditions are all related to the formation of oil at a high acidity **(Boscou, 1996)**. Saponification values varied among in the oils, and were highest in the *Pistacia lentiscus* (269,28 mg KOH/ g oil) and lowest in the *Opuntia-ficus indica* (222,3 mg KOH/g oil). The value of the *Pistacia lentiscus* oil saponification index in our study is higher than that found by Boukeloua about 193 mg KOH/g oil **(Boukeloua et al., 2012)**. The value of the saponification index of *Opuntia-ficus indica* oil of this work is higher than that oil extracted by solvent founded by El mannoubi which is 173 **(El Mannoubi et al., 2009)**, It is also higher than 186.63 for cold-pressed oil **(Gharby et al., 2013)**. The variation in the saponification index may be due to pedoclimatic factors and the stage of maturity. But generally this index is comparable with the indices of other vegetable oils such as olive, palm and avocado oil **(Karleskind, 1992)**. This shows that our studied oils extracted from the region of Ain Defla (Khemis Miliana) are rich in short chain fatty acid. the unsaponifiable fraction of *Pistacia lentiscus* oil is 1.6% and 1.8% for *Opuntia ficus indica* oil. These values are comparable to that found by Lambert for cotton oil (1,5%) **(Lambert, 2005)**,but higher than that cited by Djenotin for peanut (0,6 à 1,0) and palm oil (0,5 à 1,2) **(DjenotinN et al., 2006)**,and inferior to the one cited by Prevot for sunflower oil **(Prevot, 1987).** The peroxide values of *Pistacia lentiscus* and *Opuntia-ficus indica* oils were respectively evaluated as 6,0 and 45,8 meq/O². The value of the peroxide index obtained for lentisk oil is lower than the Codex alimentarius standard which recommends an index of 15 milliequivalents of active oxygen/kg for virgin oils and cold-pressed oils, but the value of the peroxide index found in *Opuntia-ficus indica* oil is far higher than the Standard of Codex alimentarius cited above. This very useful index informs us of conservation conditions, extraction methods, and helps us to appreciate the early stages of oxidative deterioration of the product **(Marmesat et al., 2009).** Poor storage conditions allowing oxidative degradation of triglycerides under the action of active oxygen that will attack the double bonds of unsaturated fatty acids, this allows the formation of peroxide. The low iodine value in the two oils (*Pistacia lentiscus* 44,6% and *Opuntia-ficus indica* 45,8%) may be indicative of the presence of few unsaturated bonds and would certainly contain less unsaturated fatty acids. This index allows these oils to be classified as non-siccative oils such as olive, peanut and almond oils with an iodine index set by the Codex Alimentarius standard. Based on the results obtained These two indices, index of refraction and iodine are important criteria for identifying oils.

The fatty acid composition of oil from *Pistacia lentiscus* and from *Opuntia-ficus indica* are summarised in table 02

08 fatty acids are identified in the case of *Pistacia lentiscus* oil and 11 fatty acids are identified in the case of *Opuntia-ficus indica* oil. The saturated fatty acids in the fruits oil of lentisk are palmitic, stearic, arachidic, behenic and lignoseric; however palmitic acid was the major saturated fatty acid constituent, with a percentage of 19.63%. While the acid Stearic 1% and the rest are in trace form. Concerning the unsaturated acids including palmitoleic, oleic and linoleic acids were detected in the oil studied. Oleic acid was determined to be the dominant fatty acid in the oil with a percentage of 58.35%. Linoleic acid and palmitoleic acid were detected in the oil with 19.65% and 1.15%. The saturated fatty acids in the seeds oil of *Opuntia-ficus indica* are Palmitic acid (10.169%) and stearic acid (3.581%), the others arachidic, behenic, lignocic, Pentadecanoic fatty acids appear as trace. with regard to unsaturated fatty acids are more abandoned, the main fatty acids detect are linoleic acid (63,74%), oleic acid (21,30%) as a majority fatty acids followed by palmitoleic acid, gondoic acid and erucic acid. (Tab.2)

based on these results the two oils studied are rich in unsaturated fatty acids, it is also noted that the profile of the major fatty acids of the two oils is almost the same qualitatively but quantitatively differ. *Pistacia lentiscus* fixed oil belongs to the category of monounsaturated oils, its fatty acid profile has a great similarity to olive oil and peanut oil. However, *Opuntia-ficus indica* fixed oil belongs to the category of polyunsaturated oils, its fatty acid composition is very similar to corn oil and cotton oil **(DjenotinN et al., 2006)**. The unsaturated/saturated ratio was generally high, and this high value gives these oils a good prevention of oxidation **(Charef et al., 2008)**. All this richness gives the oil of Pistacia lentiscus and *Opuntia-ficus indica* a great pharmacological cosmetologically and nutritional importance.

## 4. Conclusion

the ripe fruits of *Pistacia lentiscus* and the seeds of *Opuntia-ficus indica* provide a low level of vegetable oil, the physico-chemical parameters have shown that our products have a high saponification index and a non-negligible unsaponifiable content which gives it a high importance thanks to the noble compound it contains but *Pistacia lentiscus* oil and *Opuntia-ficus indica* oil in our case are unsuitable for use because of their high acidity and peroxide levels respectively, but the two oils represent an essential source of unsaturated fatty acid. As a result, they may offer possibilities of exploitation in the nutritional, cosmetic and therapeutic fields of agro-food technology. but they remain insufficiently poorly exploited, this opens several doors to in-depth research on its biological activities as well as the identification of the active components responsible for these activities and a better exploitation of these oils in the above-mentioned fields.

## Notes

### Competing Interest Statement

The authors have declared no competing interest.

